# Semi-automated monitoring of longitudinal microbial metabolic dynamics: A study case for lignin degradation

**DOI:** 10.1101/2024.10.18.619076

**Authors:** João Vítor Guimarães Ferreira, Gabriel Santos Arini, Nathalia Gonsales da Rosa-Garzon, Izabel Cristina Casanova Turatti, Tiago Cabral Borelli, Henrique Marcel Yudi de Oliveira Tsuji, Hamilton Cabral, Ricardo Roberto da Silva

## Abstract

This study introduces a low-cost, open-source, semi-automated bioreactor system, developed using Arduino and Raspberry Pi, for longitudinal microbial culture studies. The system protocol includes detailed instructions for custom-designed components, specifications for easily purchasable parts, and open-source code for a web-based control interface. It was validated for sterility and tested in a case study involving the cultivation of *Phanerochaete chrysosporium* and *Trichoderma reesei* to assess their metabolic dynamics, both in isolation and co-culture, for lignin degradation. Using mass spectrometry coupled with gas chromatography, several lignin degradation intermediates were annotated, including 4-hydroxybenzoic acid, vanillic acid, and ferulic acid, along with their temporal variation profiles over 25 days. The results highlighted potential synergy between the two fungal species in lignin breakdown, while also demonstrating the bioreactor’s flexibility and suitability for diverse biotechnological applications, particularly in the field of bio-degradation and waste valorization.

## INTRODUCTION

Fundamental to the carbon recycling dynamics within terrestrial ecosystems, lignin depolymerization represents not only an important technical challenge but also a focal point of interest for industries such as biorefinery (Rinaldi et al., 2016). This interest stems from the potential to derive high-value products from this intricate natural polymer, notably low molecular weight aromatic chemicals that hold promise as substitutes for petroleum derivatives (Asina et al., 2016).

Lignin, known for being a naturally occurring amorphous polymer, constitutes an essential component of lignocellulosic biomass, typically comprising 15-30% of such matrices (Grgas et al., 2023). Its complex, highly branched, three-dimensional phenolic structure comprises three primary monolignols: p-coumaryl alcohol, coniferyl alcohol, and sinapyl alcohol (Vanholme et al., 2008); while its molecular mass exhibits great variability – a consequence of the random cross-linking facilitated by phenolic group polymerization (Vanholme et al., 2008). Such polymerization involves the coupling of monolignol units via peroxidase-mediated dehydrogenation, which results in the formation of C-C bonds and aryl ether linkages, including aryl-glycerol and β-aryl ether connections (Vanholme et al., 2008).

Due to this intricate structure, lignin is highly resistant to degradation, presenting significant hurdles to its efficient utilization as a biomass resource within the industry (S. Zhang et al., 2020a). Traditionally, lignin is degraded in the pulp and paper industry using compounds such as chlorine dioxide, which are not environmentally friendly and pose additional concerns (Haq et al., 2020). In response to this challenge, the use of biodegradation has emerged as a prominent solution, offering numerous advantages, such as environmental friendliness, low energy consumption, and mild reaction condition requirements (X. Wang et al., 2017). Among the microorganisms capable of this process, white-rot fungi stand out as one of the most efficient actors, particularly due to their unique ligninolytic enzymes (Asgher et al., 2016). These enzymes possess peculiar properties, including mediator utilization and surface-active sites, which enhance their redox potential and contribute to their effectiveness in lignin degradation (Fernández-Fueyo et al., 2014). Remarkably, *Phanerochaete chrysosporium*, the most studied of all white-rot fungi, exemplifies this phenomenon through its extracellular enzymes (e.g., lignin peroxidase, manganese peroxidase, laccase), which efficiently depolymerize lignin (Mäkelä et al., 2021; X. Zhang et al., 2018).

To date, several studies have been conducted using *P. chrysosporium* and its enzymes to degrade lignin and assess compositional changes in the resultant products at the microscale. For this purpose, scanning electron microscopy (X. Zhang et al., 2019), Fourier-transform infrared spectroscopy (Halder et al., 2019), and gas chromatography coupled to mass spectrometry (Tan et al., 2022) have been commonly employed methodological approaches. However, these studies typically focus on identifying these products for a single timepoint, corresponding to a single short period of culture, failing to capture the temporal dynamics of the fungus’ behavior during the degradation process. Addressing this gap is essential, as exploring such a perspective could provide valuable insights into the changes over time in the fungus’ metabolic activities and their consequences for lignin degradation.

Another approach that has been increasingly employed to enhance the evaluation of the ligninolytic capabilities of white-rot fungi is the development of co-cultures (Luo et al., 2020). These co-cultures aim to achieve synergistic effects by combining different metabolic pathways or by inducing the expression of normally latent pathways, ultimately leading to greater efficiency in lignin degradation processes (Q. Zhang et al., 2024). One species identified as a potential contributor in co-culture with white-rot fungi is *Trichoderma reesei*, a filamentous fungus widely studied for its ability to produce cellulose and lignin-degrading enzymes (Pandya & Albert, 2014; Yan et al., 2021). Due to the extensive genetic and physiological information available on *T. reesei*, it stands out as an excellent candidate for co-culture studies (Sperandio & Filho, 2021).

To enable co-cultures over time, we introduced an automated Arduino-based bioreactor system that maintains a controlled environment and allows for periodic sampling to analyze metabolite production. The developed bioreactor system was designed to be simple, cost-effective, and easy to assemble, making it accessible to users without technical skills in electronics or programming. Its components are readily available, and its programming is user-friendly, even for those with limited experience.

This was made possible by utilizing the Arduino platform, an inexpensive, straightforward, and highly versatile microcontroller. In recent years, Arduino has become increasingly popular for laboratory applications, both in educational settings (Gerber et al., 2017) and research environments (Espeso et al., 2021). Its main advantage lies in its versatility and ease of use, enabling the creation of customized systems for a wide range of applications.

By combining these two approaches – the temporal analysis of microbial behavior in lignin degradation and the evaluation of species interactions in co-culture – this study aimed to provide new insights into lignin degradation. We monitored a co-culture of *P. chrysosporium* and *T. reesei* over a 25-day period to assess the dynamics of both their metabolite production and consumption through GC-MS analysis. This longitudinal study allowed us to observe growth variation patterns in key compounds involved in lignin degradation, such as 4-hydroxybenzoic acid, vanillic acid and ferulic acid. Additionally, the detailed documentation to reproduce a low-cost bioreactor has the potential to expand the knowledge repertoire in the temporal dynamics of lignin degradation.

## MATERIALS AND METHODS

### Development of the bioreactor

The bioreactor was built using an Arduino UNO, following the circuit design described here (https://github.com/computational-chemical-biology/arduino-based-bioreactor).

The transfers were executed using peristaltic pumps (Intllab, flow rate of 450 ml/min) with an activation duration of 15 seconds, experimentally determined based on the density of the culture medium and the tubing size so that 50 ml of medium would be transferred in each activation. A Raspberry Pi 1 B+ board was used to create a web server controlling the Arduino board, following the application design described here (https://github.com/computational-chemical-biology/arduino-based-bioreactor/tree/main/flask%20app). To improve the system, we also designed custom Teflon caps and stainless-steel cannulas at the university’s precision workshop (https://www.prefeiturarp.usp.br/page.asp?url=precisao). The description of these parts with exact dimensions are included in the project’s GitHub page.

### System validation

System validation occurred in two stages. First, a test was conducted to verify the maintenance of sterility inside the system. For this, LB medium was used in two of the glassware containers (feeding and culturing), with 200 ml and 100 ml, respectively. The set of three glassware containers and their corresponding tubing was autoclaved for 15 minutes at 115ºC. Subsequently, the system was assembled and put into operation, with the contents of the central flask (where cultivation would take place) being transferred to the collection flask, and then from the feeding flask to the cultivation flask every five days. Three complete cycles were performed. The material from the collection flask was collected every five days and discarded. At the end of the third cycle, the contents of the three flasks were plated on petri dishes with PDA medium and incubated at 30ºC for seven days.

Secondly, a monoculture of *P. chrysosporium* was carried out in a specific culture medium, with five-day cycles for a period of one month. After completion, the contents of the containers were plated on PDA medium, following the same protocol as before.

### Culture conditions

The inoculation of the cultivation unit involved using 20 discs, each 1 cm in diameter, for wild *P. chrysosporium*, and a spore suspension of *T. reesei* (QM 9414 M) at a concentration of 5.0 x 10□ spores/ml. Both species were previously cultured for 6 days at 30°C in petri dishes before their final inoculation into the system. The experiments were conducted in 5-day cycles over a period of 25 days. Each cycle began with the transfer of 50 ml from the culture flask to the collection one, followed by the transfer of 50 ml from the feeding unit to the culture one. The culture container was subjected to agitation and heating using an IKA C-MAG HS7 magnetic stirrer (agitation speed level 2, heating at 30ºC) and aeration by an aquarium air compressor (Maxxi Pro-5000). The culture medium composition was as follows: 0.7% w/v KH_2_PO_4_, 0.2% w/v K_2_HPO_4_, 0.01% w/v MgSO_4_·7H_2_0, 0.05% w/v Sodium citrate (dihydrate), 0.1% w/v Yeast extract, 0.01% w/v CaCl_2_·2H_2_O, 0.1% w/v Peptone, and 0.5% w/v Lignin, with a pH of 6.0. At the beginning, the feeding and cultivation units contained 400 ml and 100 ml of culture medium, respectively, while the collection container was empty. Throughout the experiment, the volume of the culture flask remained constant.

### Collection protocol

To ensure the maintenance of sterility within the system, samples were periodically collected from the collection flask, which was equipped with an outlet tubing sealed by a surgical clamp. On sampling days, the outlet tubing was sanitized with 70% alcohol before the surgical clamp was removed, and the contents were extracted using a sterile 60 ml syringe. The sample was then transferred to 50 ml Falcon tubes and frozen at -80ºC.

### GC-MS analysis

The analyses of the collected samples were carried out using mass spectrometry coupled with gas chromatography (GC-MS). The samples were thawed at room temperature and then centrifuged at 8,000 G for 10 minutes in order to separate de supernatant from the cellular fraction. The supernatant was collected, and the pH was adjusted to 2.0 using a 6 mol/L HCl solution. Liquid extraction was then performed with the addition of 2 mL of ethyl acetate in three separated rounds. The organic phase was collected and dehydrated with Na_2_SO4. Once dehydrated, the sample was dried under a stream of nitrogen gas and derivatized. For derivatization, 20 μl of pyridine and 100 μl of bis (trimethylsilyl)trifluoroacetamide (BSTFA) were added. The solution was placed in a water bath at 60ºC for 30 minutes, with regular shaking. The resulting silylated samples were then injected into GC-MS QP2010 Ultra (Shimadzu, Kyoto, Japan). A ZB-5MS capillary column (30 m × 0.25 mm inner diameter × 0.25 µm film thickness) was used, with helium as the carrier gas at a flow rate of 1 ml/min. The column temperature started at 50ºC (held for 5 minutes) and was increased to 300ºC at a rate of 10ºC per minute. The injection temperature was 280ºC, with the ion source maintained at 250ºC and the transfer line at 200ºC. The solvent delay was set to 5 minutes. The injection volume was 1 μl.

### Data processing

The Shimadzu QGD files were converted to an open format (.mzXML) with OpenChrom software (v. 15.0) (Wenig & Odermatt, 2010) (https://lablicate.com/platform/openchrom) (accessed in February 2024) and pre-processed through MZmine 2.53 (Schmid et al., 2023) following the parameters presented in the .xml file available in the project’s GitHub page. The pre-processed data were subsequently annotated through spectral pairing through the GNPS platform (M. Wang et al., 2016), using the MOLECULAR-LIBRARY SEARCH-GC workflow. The full results of the spectral matching, as well as all the adopted parameters, are available at https://gnps.ucsd.edu/ProteoSAFe/status.jsp?task=cc5af8857436441f9373a6bda0a17199 (accessed on 7 July 2024).

### Statistical Data Analysis

To verify the correlation between the peak areas for each compound annotated at the evaluated timepoints, an analysis was conducted using the Pearson correlation coefficient. A threshold value of 0.80 (in absolute terms) was established to filter correlations between the peak areas under study. The detailed calculations and analysis steps can be reviewed in the Jupyter Notebook available in the project’s GitHub page.

## RESULTS AND DISCUSSION

### Design X development of the bioreactor

A schematic of the system can be seen in Fig. 1a-b. In simplified terms, the developed bioreactor system consists of three essential parts:

**Figure 1.**
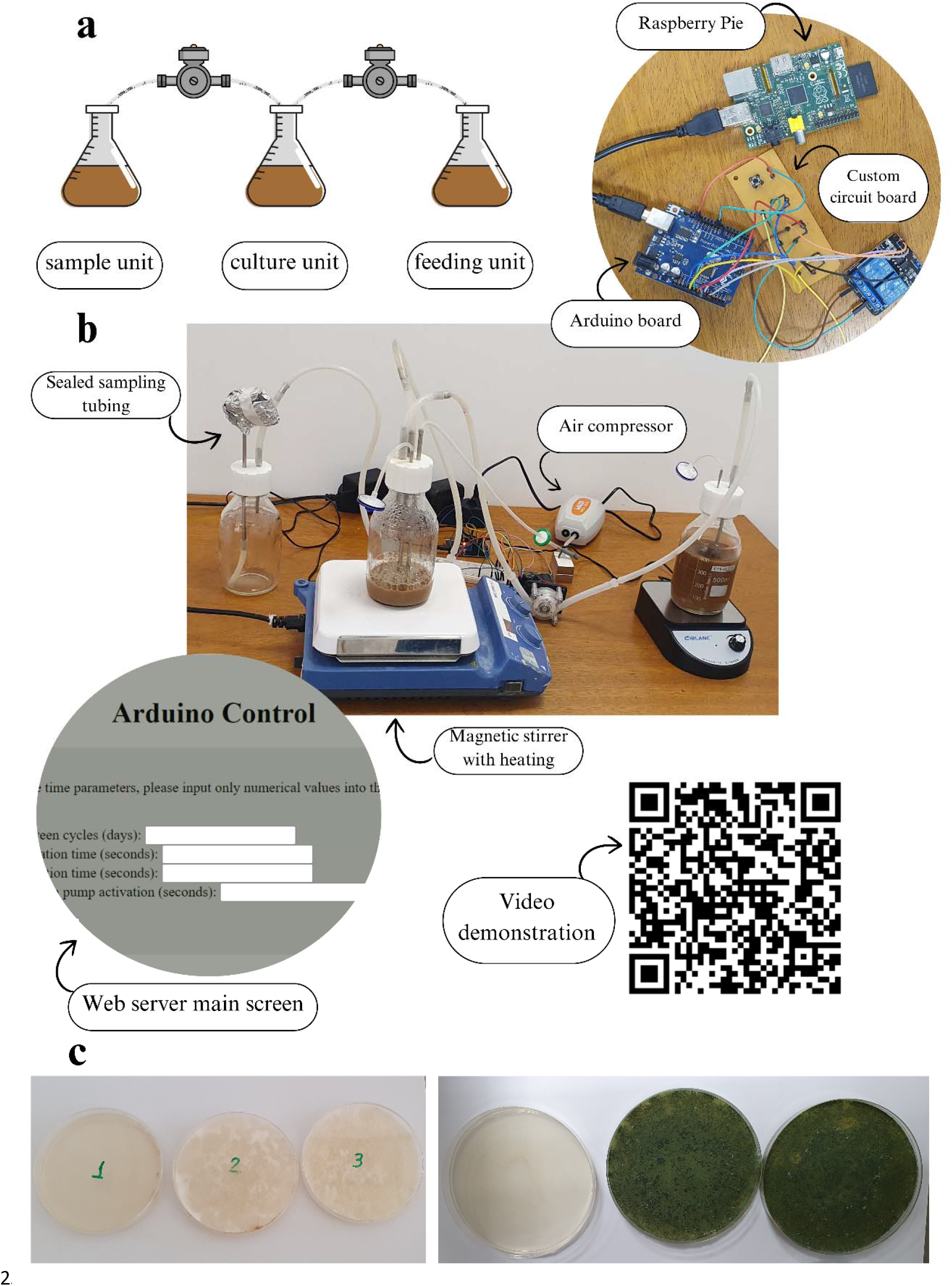
Overview of the bioreactor system and post-cultivation contamination analysis. A) Schematic diagram of the bioreactor setup. B) Photograph of the operational system, highlighting key components, including the customized circuit, Arduino and Raspberry Pi control boards, and the web server interface. A video demonstration of the system in action is accessible via the QR code. C) Petri dishes from the *P. chrysosporium* (left) and *T. reesei* (right) cultivation cycles, demonstrating the system’s effectiveness in preventing external contamination. For each cycle, the three plates represent samples from the feeding unit (left, which remained sterile), the cultivation unit (center), and the collection unit (right).

1. Three 500-ml glass containers serving as the feeding, cultivation, and collection units of the system, equipped with custom-made Teflon caps with spaces for attaching stainless-steel cannulas through which the contents of the containers pass.
2. Two peristaltic pumps, connected to the stainless-steel cannulas via silicone tubings, directing the transit of the contents.
3. A central control system, consisting of an Arduino UNO board, an integration circuit board, and a Raspberry Pi 1 B+ board, responsible for storing and executing the programmable commands that govern the overall functioning of the system.

Additional elements that facilitated the operation of the experiments included magnetic stirrers with heating, an air compressor for aquariums, 0.22 μm membrane filters, and a surgical clamp. The magnetic stirrers maintained the culture temperature and ensured uniform distribution of nutrients in the culture medium. The air compressor aerated the cultivation medium, which was essential for the growth of the species studied. The membrane filters prevented pressure imbalances and system contamination during the experiments. Similarly, the surgical clamp prevented contamination by the outlet tubing, which was only opened during sample extraction, in accordance with a validated protocol.

#### Arduino e Raspberry Pi digital controllers

an Arduino UNO microcontroller was used to control the activation of the peristaltic pumps and, consequently, the movement of the contents of each flask at defined intervals. The flow rate of the pumps was measured beforehand, and the required activation duration of each pump was estimated and programmed into the external software (Arduino IDE) used to control the system. The connection to the Raspberry Pi provided internet access to the Arduino, allowing changes to the duration of pump activation and the size of the cultivation cycle, as well as monitoring of all stages of the process, to be carried out via a web application developed specifically for this purpose.

#### System Validation

The assembly was validated through sterility tests, confirming the absence of external contamination within the system. In the first test, conducted with LB medium for three cycles and incubated at 30ºC, no signs of contamination were detected. In the second test, performed at the end of five cycles with *P. chrysosporium*, only *P. chrysosporium* was found in the cultivation and collection flask samples, while no microorganisms were detected in the feeding container samples (Fig. 1c). This result indicates not only the protection of the medium against external contamination but also the sterile integrity of the feeding unit.

### Lignin degradation intermediates

Under extraction with ethyl acetate, GC-MS analysis of the collected samples identified degradation intermediate products consistent with those reported in the literature (Duan et al., 2016; Tan et al., 2022). Four main groups of compounds were observed, including organic acids (e.g., lactic acid, 2-hydroxybutanoic acid, 4-hydroxybenzoic acid, palmitic acid, stearic acid), esters (e.g., bis (2-ethylhexyl) phthalate), aromatic hydrocarbons (e.g., m-xylene, phenol), and alcohols (e.g., 2-ethoxyethanol, ethylene glycol).

Lignin degradation occurs in two main stages (Tan et al., 2022). Initially, depolymerization of lignin polymers is marked by the oxidation of lignin in the presence of air, involving the extraction of an electron from lignin, the consequent formation of free radicals from the substrate molecules, and the reduction of oxygen to water (S. Zhang et al., 2020b). Lignin is then degraded through catalytic polymerization and depolymerization reactions by enzymes such as laccase, caused by the instability of the formed free radicals. Furthermore, non-enzymatic reactions induced by the abundance of free radicals lead to decarboxylation, demethoxylation, and carbon-carbon chain breaks (S. Zhang et al., 2020b). The lignin polymer subsequently converts into oligomers that proceed to the second phase of degradation, where they are metabolized into smaller benzene-derived molecules (Tan et al., 2022). At this stage, the resulting free radicals replace hydrogen atoms in the side chains of aromatic compounds, forming alcohols. Consequently, a large number of aromatic alcohols is produced during lignin degradation.

In our analysis, the presence of several small aromatic compounds was noted, such as dimethyl phthalate, 4-hydroxybenzoic acid, benzoic acid and 3-tert-Butylphenol – all four described in lignin degradation processes.

Dimethyl phthalate, like many phthalate derivates, is an intermediate that, in later stages of degradation, generates phthalic acid, which in turn is metabolized into syringyl, protocatechuic acid, and other phenolic compounds, which are further degraded (Tan et al., 2022). In the temporal analysis, a pronounced reduction of this compound was observed over time in both the *T. reesei* monoculture and the co-culture (Fig. 2b).

**Figure 2.**
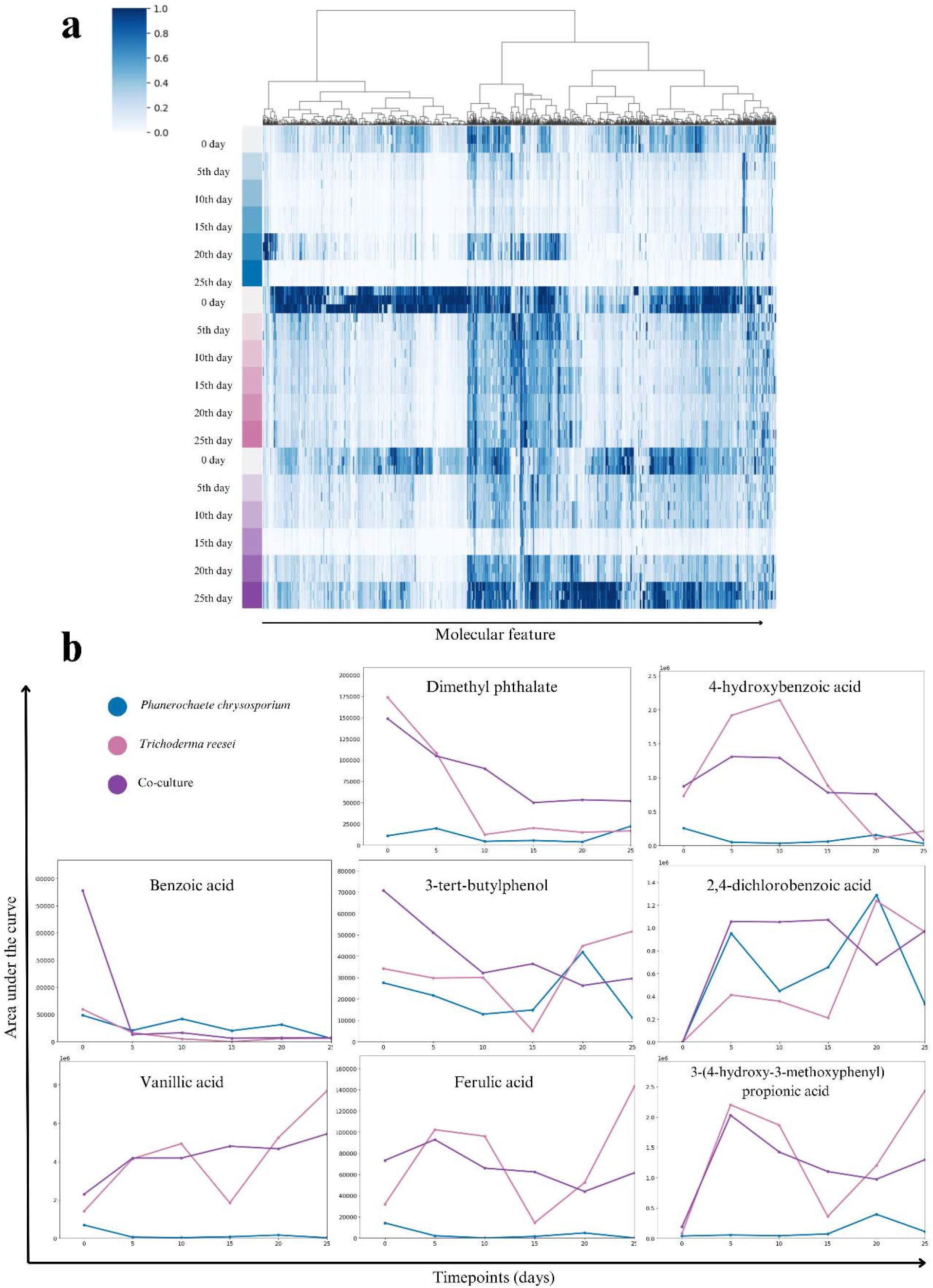
Monitoring of compound levels over a 25-day period. A) Heatmap depicting the concentration of compounds detected across three cultivation groups: *P. chrysosporium* monoculture, *T. reesei* monoculture, and co-culture of the two species. In the heatmap, blue shades indicate the area under the peak, with darker shades representing higher concentrations. B) Graphs showing the growth and decline patterns of the area under the curve for key compounds involved in lignin degradation.

Benzoic acid and its derivate 4-hydroxybenzoic acid are central intermediate in several lignin degradation pathways, such as the gentisic acid pathway, coumaric acid pathway, and cinnamic acid pathway (Pérez-Pantoja et al., 2008). In the temporal analysis, a diminution in the area of benzoic acid peaks was observed across all three culture conditions compared to the initial condition (timepoint 0 - medium without culture), with such decrement being most distinct in the *T. reesei* monoculture (Fig. 2b). Conversely, 4-hydroxybenzoic acid levels increased in both the *T. reesei* monoculture and the co-culture. This increase is likely attributable to the activity of benzoate 4-monooxygenase, an enzyme present in the *T. reesei* genome (Grossetête et al., 2010) that can convert benzoic acid into 4-hydroxybenzoic acid.

Tert-butylphenol derivatives have been the focus of recent studies that demonstrate their involvement in microbial lignin degradation pathways, such as the resorcinol pathway (Toyama et al., 2010). Considering the variable behavior of this molecule over time, with a more notable decrease observed in the co-culture, it can be suggested that *P. chrysosporium, T. reesei*, or their combination may possess a similar pathway, where the tert-butyl structure is retained and metabolized via the meta-cleavage pathway (Fig. 2b).

#### Chlorinated compounds

Due to the lignin being sourced from pulp and paper industry waste, a significant number of chlorinated compounds were detected in the analyses. Among these, 2,4-dichlorobenzoic acid was particularly noteworthy, exhibiting rapid accumulation during the first five days and subsequently stabilizing across the three analysis groups (Fig. 2b). This could be attributed to the chlorination of the lignin polymeric structure, a process commonly employed in the paper and cellulose industries (Haq et al., 2020).

This benzoic acid derivative likely resulted from reactions similar to those that produced compounds such as 4-hydroxybenzoic acid, which also showed an increase during the first 10 days in the *T. reesei* monoculture and co-culture. However, unlike 2,4-dichlorobenzoic acid, 4-hydroxybenzoic acid did not remain stable, instead experiencing a decline over the last 15 days (Fig. 2b). This suggests that 2,4-dichlorobenzoic acid may follow a different metabolic pathway than 4-hydroxybenzoic acid or may be more resistant to degradation due to chlorination.

#### Co-culture outcomes

Upon analyzing the annotated compounds that exhibited an increase or decrease in the area under the curve across the observed time points, it was found that the co-culture generated a greater number of compounds with both positive and negative correlations (116 compounds) compared to the isolated cultures (91 compounds for *P. chrysosporium* and 35 compounds for *T. reesei*). Only two compounds displayed similar behavior between the monocultures. Sixteen compounds were common between *P. chrysosporium* and the co-culture, while seven were common between *T. reesei* and the co-culture. Notably, no compound exhibited a significant correlation across all three groups.

Considering only the compounds reported in the literature as lignin degradation products (Ebrahimi et al., 2024; Xia & Stone, 2019; Zhu et al., 2018), the co-culture yielded 11 compounds with significant correlation, including guaiacol, adipic acid, vanillic acid, and malonic acid. In comparison, *P. chrysosporium* and *T. reesei* each produced 4 such compounds. Among these, 2-methoxy-5-methylphenol was common to both the *P. chrysosporium* monoculture and the co-culture, while adipic acid was found in both the *T. reesei* monoculture and the co-culture.

The analysis reveals a diverse metabolic response within the co-culture, as indicated by both the increased number of compounds with significant correlations and their greater diversity. This suggests a potential synergistic interaction between the two species, particularly in relation to lignin degradation. Such interactions likely lead to the development of unique and distinct metabolic pathways compared to those observed in the monoculture.

In terms of exclusivity, three annotated compounds (N-acetyl-leucine, phosphine, 1,3-propanediylbis[dicyclohexyl-], and 1,3-dithiaindane, 2- (3-methylbutoxy)-4- (trifluoromethyl)) were detected in both the *T. reesei* monoculture and the co-culture, but were absent in the *P. chrysosporium* monoculture. Notably, no compounds were found to be exclusive to either *P. chrysosporium, T. reesei*, or the co-culture alone. This absence of exclusive compounds across the conditions suggests a shared metabolic environment, where certain metabolites are either consistently produced or inhibited depending on the presence of specific interactions between the species.

When focusing on compounds typically associated with the final stages of lignin degradation (Ebrahimi et al., 2024; Xia & Stone, 2019; Zhu et al., 2018), as outlined in Fig. 3, evidence of synergism between the species becomes more apparent.

**Figure 3.**
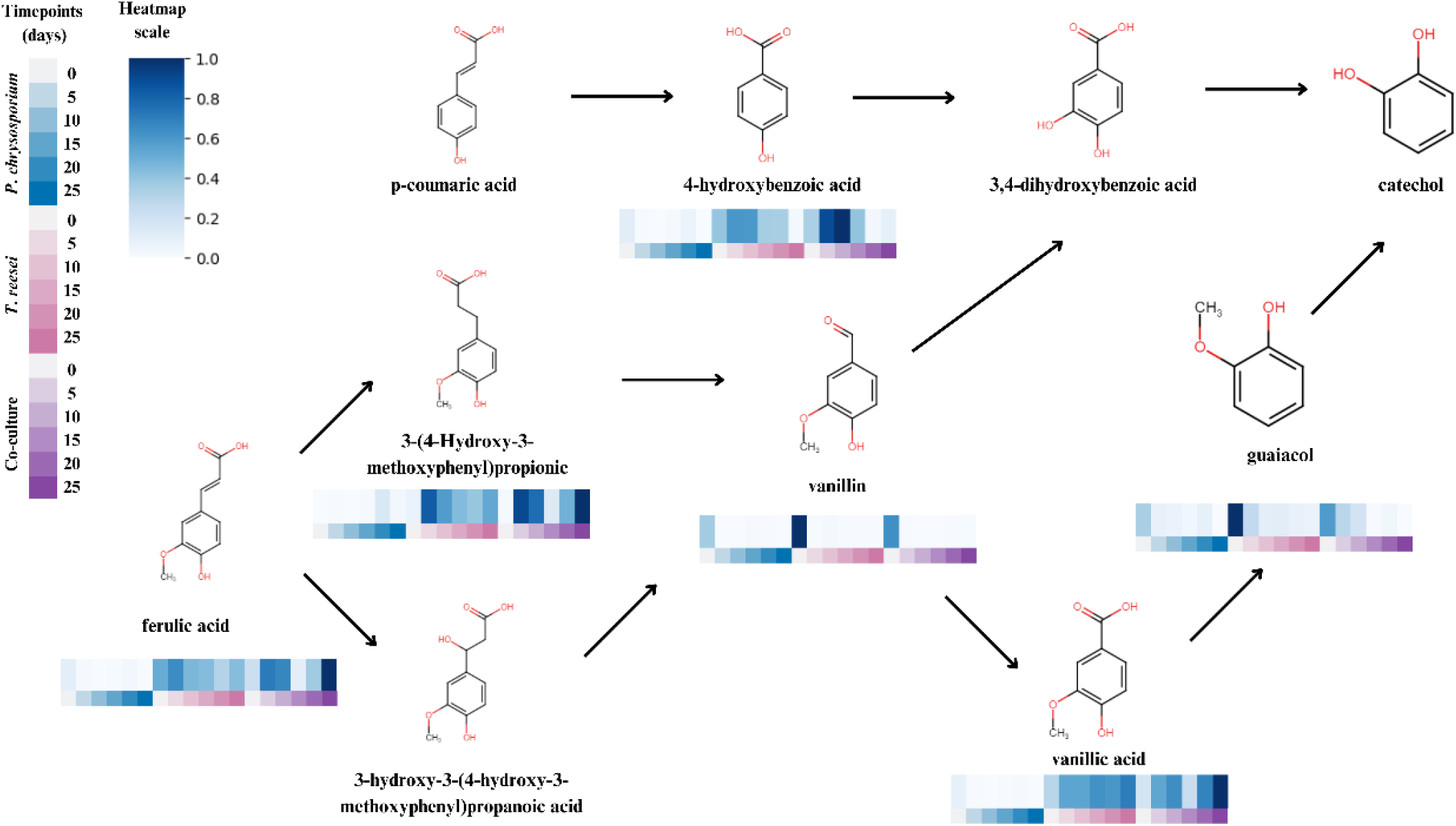
Proposed final stages of lignin degradation, highlighting the identified compounds within the system. The accompanying heatmap illustrates the variation in the areas under the curve for these compounds across different timepoints.

For instance, with regard to vanillic acid, the area under the curve over time shows that in the *P. chrysosporium* monoculture, vanillic acid levels remain very low, indicating its continuous consumption. In contrast, in the *T. reesei* monoculture, these levels consistently increase, suggesting ongoing production. In the co-culture, however, a balance is observed, implying that the consumption by *P. chrysosporium* and the production by *T. reesei* are effectively counterbalancing each other over time (Fig. 2b). This balance reflects a dynamic interaction between the species, likely contributing to a more efficient and sustained metabolic process in the co-culture.

Similarly, ferulic acid and 3- (4-hydroxy-3-methoxyphenyl)propionic acid were maintained at generally lower levels during co-culture compared to the *T. reesei* monoculture, where these compounds appeared to increase towards the end of the observation period (Fig. 2b). This reduction in levels during co-culture suggests that the interaction between the species may influence the metabolic fate of these compounds, possibly through enhanced consumption or altered metabolic pathways, resulting in a distinct profile from that of the monocultures.

## CONCLUSION

Overall, the results of this investigation suggest a synergistic interaction between *P. chrysosporium* and *T. reesei* in lignin degradation. However, further studies are needed to deepen our understanding of this interaction and to explore ways to optimize it. Genomic analyses, in particular, could provide valuable insights into the molecular mechanisms underlying this synergy and identify potential targets for enhancing ligninolytic activity. Additionally, investigations that explore variations in environmental conditions or cultivation periods may offer strategies to increase enzyme production and enhance the lignin degradation capabilities of these species. Our open source bioreactor project can be expanded to test a wide range of culture conditions, offering new insights into the metabolic dynamics. Such research could contribute significantly to the development of more efficient biotechnological applications for lignin valorization.

## ACKNOWLEDGEMENTS

This work was supported by Fundação de Amparo à Pesquisa do Estado de São Paulo (award numbers 2017/18922-2; 2020/02207-5; 2021/10401-9; 2022/15042-0).

